# Corn rust population genomics reveals a cryptic virulent group and adaptive effectorome

**DOI:** 10.1101/2025.05.07.652763

**Authors:** Yuanjie Li, Peng Zhao, Xiufeng Liu, Clement K.M. Tsui, Daniel Croll, Junmin Liang, Lei Cai

## Abstract

Understanding the population structure of pathogens and the genetic determinants driving virulence gains is crucial for managing epidemic plant diseases. *Puccinia polysora*, a giga-scale fungal pathogen causing southern corn rust, has posed significant threat to global food security recently. Traditionally, *P. polysora* was considered clonal with minimal genetic variation. However, our population genomic and transcriptomic studies conducted in China, the emerging epicentre of the disease, have challenged this view. By adopting variant analyses appropriate to the dikaryotic nature of the pathogen, we discovered an unexpectedly clear population structure with six distinct groups. A cryptic group exhibits high virulence, facilitated by group-specific variation, and diversification of effectors. Although the Chinese population of *P. polysora* is predominantly asexual, internuclear exchange on some chromosomes have introduced recombination signals. The comprehensive pan-effectorome analyses revealed substantial presence/absence variation and alternative splicing events on effectors, shaping a highly adaptive effector repertoire in *P. polysora*. In conclusion, our findings highlight the tandem mapping on exploring clear genetic structure of dikaryotic species and reported the emergence of a virulent group and an adaptive effectorome of *P. polysora*. Effective containment strategies must be flexible to counter the threats posed by the unexpectedly dynamic evolution of this pathogen.

## Introduction

The emergence and dissemination of novel pathogens or pathogen races pose a significant threat to global food security [1]. Particularly, fungal pathogens are responsible for approximately 10-80% of crop losses [2,3]. There has been a noted rise in the occurrence of fungal diseases in agricultural systems, including wheat blast [4], wheat stem rust by the Ug99 group [5] and Dutch elm disease [6]. In order to effectively control fungal pathogens and implement robust disease management strategies, it is essential to gain a comprehensive understanding of their population structure, the genomic determinants that shape their adaptations as well as the ecological and evolutionary processes that act upon pathogen populations.

*Puccinia polysora* f.sp. *zeae* (*Ppz*) (Pucciniales, Basidiomycota) is a biotrophic fungus that causes southern corn rust (SCR) (Fig. 1a), one of the top ten most prevalent diseases on maize [3]. Historical records show widespread SCR epidemics in the tropics, including South Africa [7], Southern USA [8,9] and Southeast Asia [10,11], causing up to 50% maize yield loss. In recent decades, SCR has been observed spreading to the temperate zones, particularly in China [12,13] and the USA [14–16], posing increasing risk on global maize production (Ramirez-Cabral et al., 2017). Given the frequent outbreaks in China, the Chinese government has listed SCR as one of top-priority crop diseases (Announcement of the Ministry of Agriculture and Rural Development of the People’s Republic of China No. 654, 2023), reflecting the urgency and importance of addressing this threat to Chinese crop production. Until now, studies on *P. polysora* population using molecular markers are very limited [11,17], and the genomic determinants shaping virulence variation and adaptive evolution remain poorly understood.

**Fig. 1.**
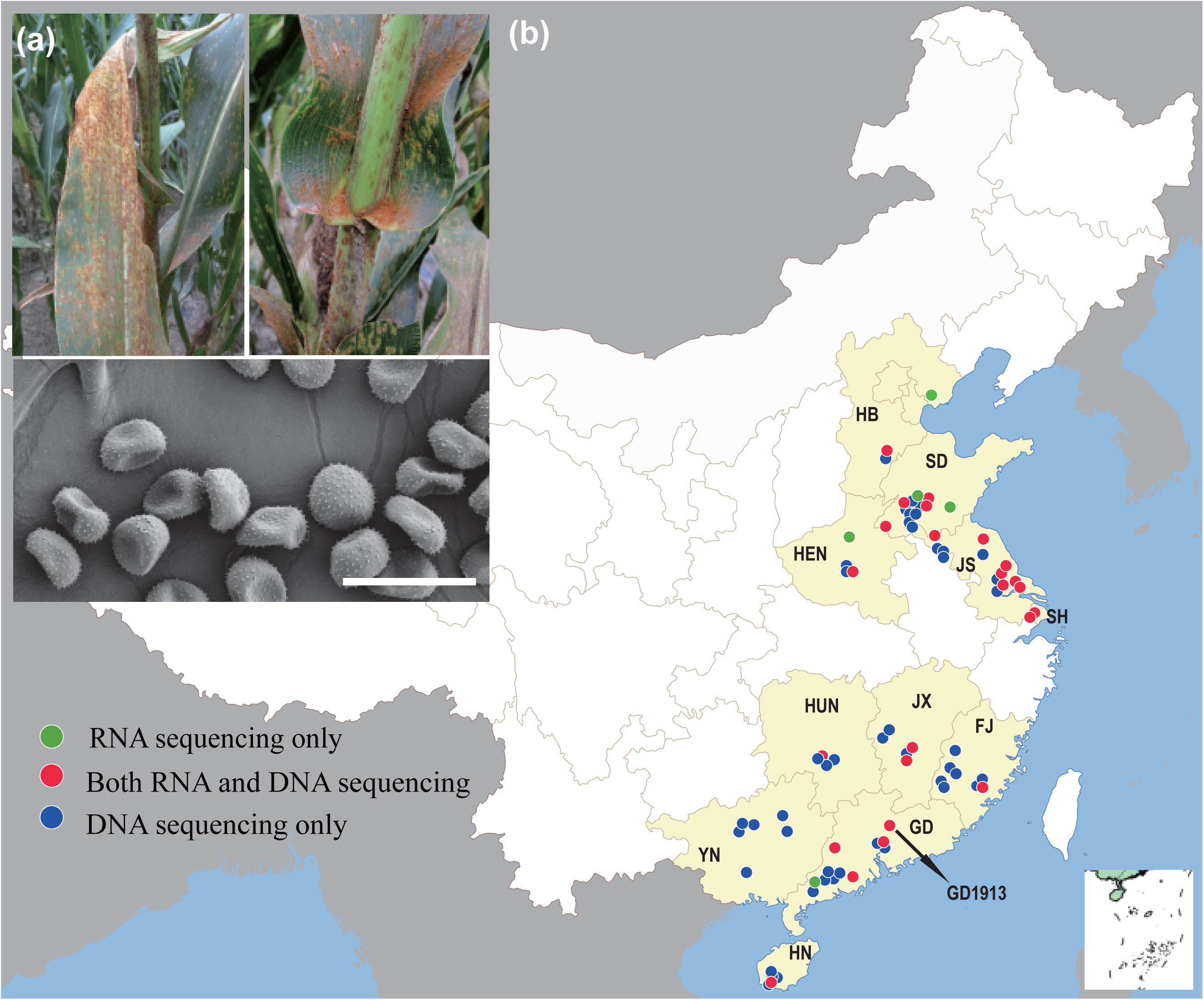
| Symptom of southern corn rust and the sample types and distributions. (a) Disease symptoms and urediniospores on corn leaf. Bar = 50 _μ_m. (b) Sequencing type and distribution of samples analysed in this study. The location of GD1913, referred to as the reference genome is indicated.

Population genomics investigations have yield valuable insights into the ecological and evolutionary processes of fungal plant pathogens [18]. However, unique biological characteristics of *Puccinia* species, such as their biotrophic parasites, complex life cycles dominated by dikaryotic spores, large genome size and high repeat contents [19,20], have hindered the accessibility of complete genomes of *Puccinia* spp. [21–23]. Until recently, Hi-C technology enabled the haplotyping of genomes from a few rust species, including *P. polysora* [24–28]. Meanwhile, a series of questions raised for dikaryotic fungi. The performance and impact of mapping reads to individual haplotypes versus both haplotypes on population structure remain to be fully elucidated. Previous studies have suggested that the Chinese *P. polysora* population exhibited a weak population structure, as inferred by nine simple sequence repeat (SSR) markers [17] and SNPs mapped to a single haplotype [27]. However, it cannot been ignored that as obligate biotrophs, *Puccinia* spp. can evolve rapidly under the selective pressures exerted by their host plants [29,30]. Since the 1990s, China has experienced a sustained epidemic of SCR for at least 30 years, accompanied by several rounds of cultivar turnover [12,31]. Therefore, it is highly likely that *P. polysora* populations in China are subject to differentiation by host selection. Whether the observed weak genetic structure of *P. polysora* is an artifact resulting from the use of a single haplotype reference genome warrants further exploration.

Understanding the driving force for virulence evolution is critical for the management of plant pathogens. The life cycle of rusts is complex, which has a major impact on the virulence evolution [32,33]. Sexual reproduction provides rusts with evolutionary innovations to overcome plant resistance, allowing for rapid adaptation [34–36], while asexual reproduction preserves highly adapted genotypes, facilitating disease outbreaks and epidemics. Meanwhile, Figueroa et al. [32] proposed three asexual processes associated with the evolution of virulence in rust fungi: mutation, internuclear exchange and somatic fusion and exchange. Recent studies have highlighted the significant contribution of somatic hybridization in generating genetic diversity within rust populations [24,28,37]. *P. polysora* is currently known only for its asexual stage with infinite infection by urediniospores. Because the sex-specific precursor, telia of *P. polysora*, is extremely rare in nature and has never been found under laboratory conditions [38,39], *P. polysora* is tentatively considered to be an asexual reproduction population. The germination condition of teliospores of rust fungi is usually elusive, that make the sexual period difficult to be detected in nature [40]. It took a century for *P. striiformis* (wheat rust pathogen), which had been thought to reproduce asexually, to report a sexual cycle [41]. By detecting the mating loci, Holden et al. [42] reported a positive relationship between genotypic diversity at mating-type loci and the potential ability for sexual reproduction in *P. striiformis* populations. It will be a convenient method to assess the sexual reproductive potential of rust population without lengthy inducing germination of teliospores. Our study will investigate the genotypic diversity of mating loci, and evaluate the roles of internuclear exchange, somatic or potential sexual recombination in the genetic differentiation of *P. polysora*.

Effectors, one of the key virulence factors of plant pathogens, play a pivotal role in establishing successful infection [43,44]. The Pan-effectorome represents the diversity of effectors at the population scale, reflecting the adaptability and variability of pathogens in countering host immune responses. Deciphering this landscape is important for disease control through resistance breeding. Host-recognised effectors, known as avirulence factors, are increasingly employed to expedite the deployment of resistance genes [43,45], while core effectors contribute to the development of broad-spectrum resistance breeding [46]. The reproductive strategy of a pathogen population can significantly influence the openness or closeness of its pan-effectorome. Those that reproduce predominantly by clonal means tend to exhibit a fairly homogeneous genome structure within their lineages [47,48], resulting in lower diversity of effector repertoire. These populations often exhibit smaller effective population sizes, which correlate with increased clonality [49]. Therefore, a closed pan-effectorome would be expected if *P. polysora* is clonal. Pan-effectorome studies in rust fungi are rare. Recently, the presence/absence variation (PAV), particularly in effector genes, has been highlighted to significantly affect the virulence spectra of plant pathogens and the strain-specific dispensable regions, usually harbouring pathogenicity-related genes, are crucial to study virulence evolution [50–52]. Therefore, the virulence plasticity of *P. polysora* may be underestimated by analyses based on a single reference genome [27], necessitating the adoption of a pan-effectorome approach.

To address the above issues, we performed a comprehensive population genomic and transcriptomic analyses of *P. polysora* using 76 DNA resequencing and 33 RNA-seq datasets. We adopted a tandem mapping strategy, using two phased haplotypes together as the reference genome to minimise inter-nuclear variant noise. We revisited the resequencing data from Liang et al. [27], which was initially mapped to a single haplotype. Our results supported clear genetic structure with significant differentiation, in contrast to the weak population divergence previously reported [17,27]. The inoculation assay across five resistant inbred lines revealed a highly virulent group characterised by specific variation in effectors and mating-type loci. By incorporating transcriptome data from infected plant tissues, we were able to construct the first pan-effectorome landscape for *P. polysora*. Our results suggest that to complement the negative effects of clonal reproduction, *P*. *polysora* may adopt a loose TEs defence mechanism and internuclear exchange events to enhance genetic variation. Therefore, *P. polysora* is equipped with an open pan-effectorome that exhibits a high degree of virulence variation through presence/absence variation and extensive alternate splicing in effectors. Our work not only provides a valuable resource for future research on plant-microbe interactions and resistance breeding, but also improves our understanding of the adaptive evolution of *P. polysora*, contributing to the development of effective disease management and control strategies.

## Results

### Tandem mapping excludes inter-nuclear variation

We compared two mapping strategies: a haplotype-specific mapping approach with a single phased genome (either hapA or hapB), and a tandem mapping approach utilizing two haplotypes as the reference genome (hapA+hapB). The two mapping strategies obtained 5,474,855 (hapA) and 81,384 (hapA+hapB) SNPs, respectively. The use of tandem mapping strategy resulted in a significant decrease in the number of SNPs (Fig. 2a, Fig. S1). The phylogenetic tree conducted using hapA+hapB revealed shorter branch lengths between isolates within the same group, but highlighted a more pronounced differentiation level between different groups (Fig. 2a). For example, the genetic distance between the bottom group (marked in purple) and its adjacent group (marked in blue) is approximately 28 times greater when using the hapA+hapB method compared to the tree constructed based on SNPs mapped to hapA alone (Fig. 2a). Taking the *AvrRppC* gene as an example, four SNPs were detected when analysed using either the hapA or hapB approach individually. However, two of these SNPs (SNP 1 and 2) were actually inter-nuclear variants (Fig. 2b, c) and no longer appeared as heterozygous when assigned to distinct haplotypes in the tandem mapping mode. The remaining two SNPs (SNP 3 and 4) represent true inter-individual genetic variation (Fig. 2c). Additionally, the SNP read depth when mapping to single haplotype (hapA or hapB) is twice (60 ×) that of mapping to both haplotypes (30 ×), indicating reads correctly assigned to two haplotypes.

**Fig. 2.**
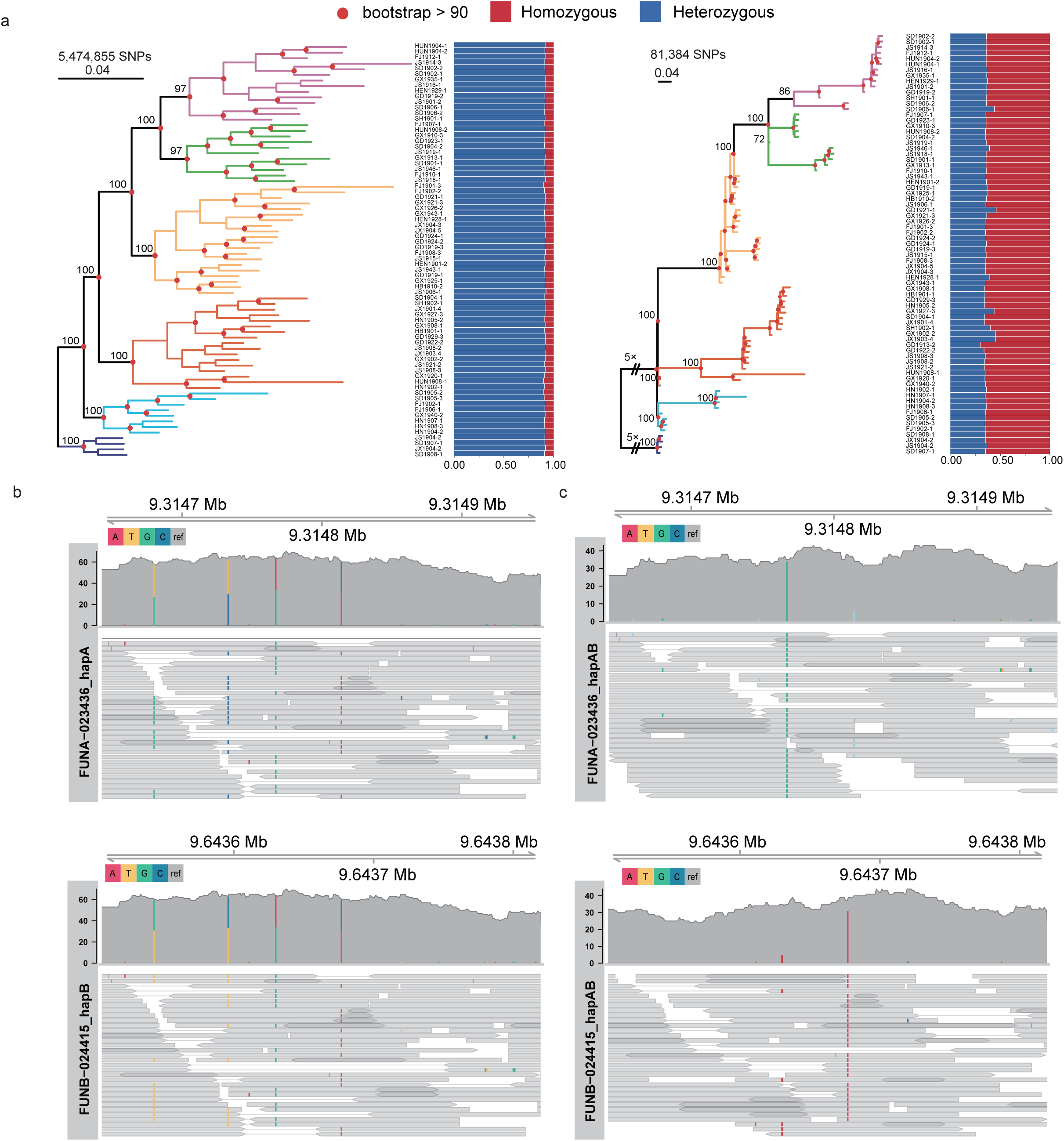
| Comparisons of two mapping strategies. (a) The phylogenetic tree constructed using SNPs by mapping short reads to hapA (left) and hapA+hapB (right). The bar charts right to the trees show the percentage of heterozygous and homozygous sites for each isolate. Six major clades are colored separately. (b) The representative isolate SD1908-1, was analysed for its reads mapping plot by using a single haplotype (either hapA or hapB) as the reference genome. The *AvrRppC* was utilized as a case to illustrate the variations in SNPs observed in the two distinct mapping approaches. The mapping regions are Ppz_chr14A: 9,314,643-9,314,957 (FUNA_023436, *AvrRppC* in hapA) and Ppz_chr14B: 9,643,506-9,643,820 (FUNB_024415, *AvrRppC* in hapB). (c) The representative isolate SD1908-1, was analysed for its reads mapping plot by using hapA+hapB as the reference genome.

### Population structure and the emergence of a highly virulent group

Utilizing the variants obtained from the tandem mapping mode, we reassessed the population divergence of Chinese *P. polysora* using discriminant analysis of principal components (DAPC). The optimal population number (*K*) was detected as either 6 or 7, with the lowest BIC values (Fig. S1d). We chose 6 as best *K* where the populations exhibited clear demarcations consistent with the phylogenetic clades determined by the IQ-tree. The average nucleotide diversity (π) of the *P. polysora* population was 1.98 × 10^-5^, with G6 recording the highest (2.38 × 10^-5^), and G3 the lowest (1.77 × 10^-5^) (Fig. S1c). In the principal component analysis (PCA), the top three principal components explained 65% of the genetic variance observed (Fig. 3b). The G6 group was notably distinct from the rest, characterised by high *Fst* values (> 0.25) between G6-containing pairs (Fig. 3c). Group differentiation did not show a correlation with the geographical regions, because several groups contain samples from different regions (Northern China, Southern China and Central China) (Fig. 3a). However, the virulence differentiation between G6 and other groups are significant. The inoculation assay (two representatives randomly selected from each group) demonstrated that the G6 strains were more virulent (disease index > 5) against the main resistant inbred lines compared to other groups (disease index < 3) (Fig. 3d, Fig. S2).

**Fig. 3.**
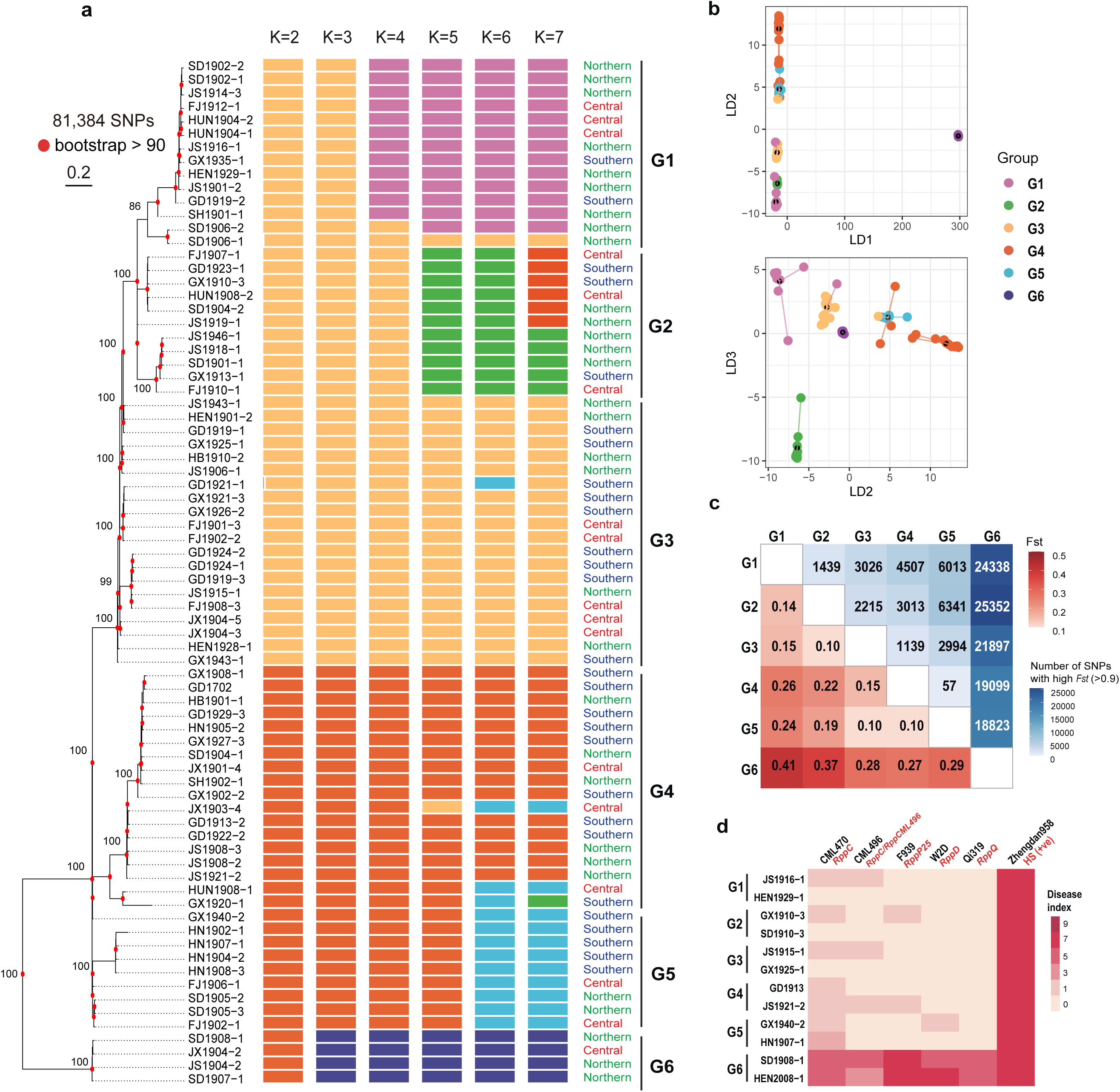
| Statistics of variants and population structure. (a) Composition proportions inferred with *K* ranging from 2 to 7. The maximum likelihood tree was constructed using 81,384 SNPs and highly supported nodes (with bootstrap values ML > 90) are labeled in red dots. (b) The DAPC analyses plotted by top three LDs. Black dots represented central of each genetic group. Eigenvalues of LDs are shown in Fig. S1e (c) Pair-wise *F*st (above diagonal) and number of high *F*st (> 0.9) SNPs (below diagonal) in all groups. (d) Inoculation assay of 12 representative isolates on five inbred lines with known resistant genes. The high susceptible variety “Zhengdan958” was used as positive control. The disease index was recorded mean value from three distinct biological replicates. The values with no spore lesion were denoted by “0”, while the presence of spore lesions covering 1%-5%, 5%-25%, 26%-50%, 51%-75%, and 76%-100% of the area were scored as “1”, “3”, “5”, “7” and “9”, respectively. The inoculation symptoms are shown in Fig. S2.

High *F*st is indicative of diversifying selection. We further investigated the group-specific genetic variation using an outlier approach–SNPs with high *F*st values (>0.9) among six groups. A total of 28,873 SNPs were identified, of which 64.38% were unique to G6, distinguishing it from other groups. The majority (98%) of the SNPs were intergenic, with minimal impact on protein structure (Table S1). SnpEff annotation revealed that 272 variations had moderate or strong functional effects (Table S1). Most SNPs were located in candidate genes of unknown function. The 37 nonsynonymous SNPs detected in genes involved in regulation, nutrition & metabolism, and pathogenicity & adaptation showed group-specific variation (Table S2, Fig. 4). Specifically, SNPs in G6 were enriched in secreted effectors, such as FUNA_023436-T1/FUNB_024415-T1 (*AvrRppC*), FUNA_005021-T1, FUNB_009431-T1 and FUNA_017151-T1 (Fig. 4). We then detected GO functional annotations for G6-specific variants (Table S3). Among 113 identified terms, transport-related terms, e.g., vesicle-mediated transport (GO:0016192), peptide transport (GO:0015833) and intracellular transport (GO:0046907), are detected in multiple genes (Godinho et al., 2014). In addition, protein modification process, such as GO:0036211 and GO:0018193, were also involved. However, only limited genes have annotations, no GO terms were significantly enriched GO terms (*P* > 0.05 in each GO term) (Table S3).

**Fig. 4.**
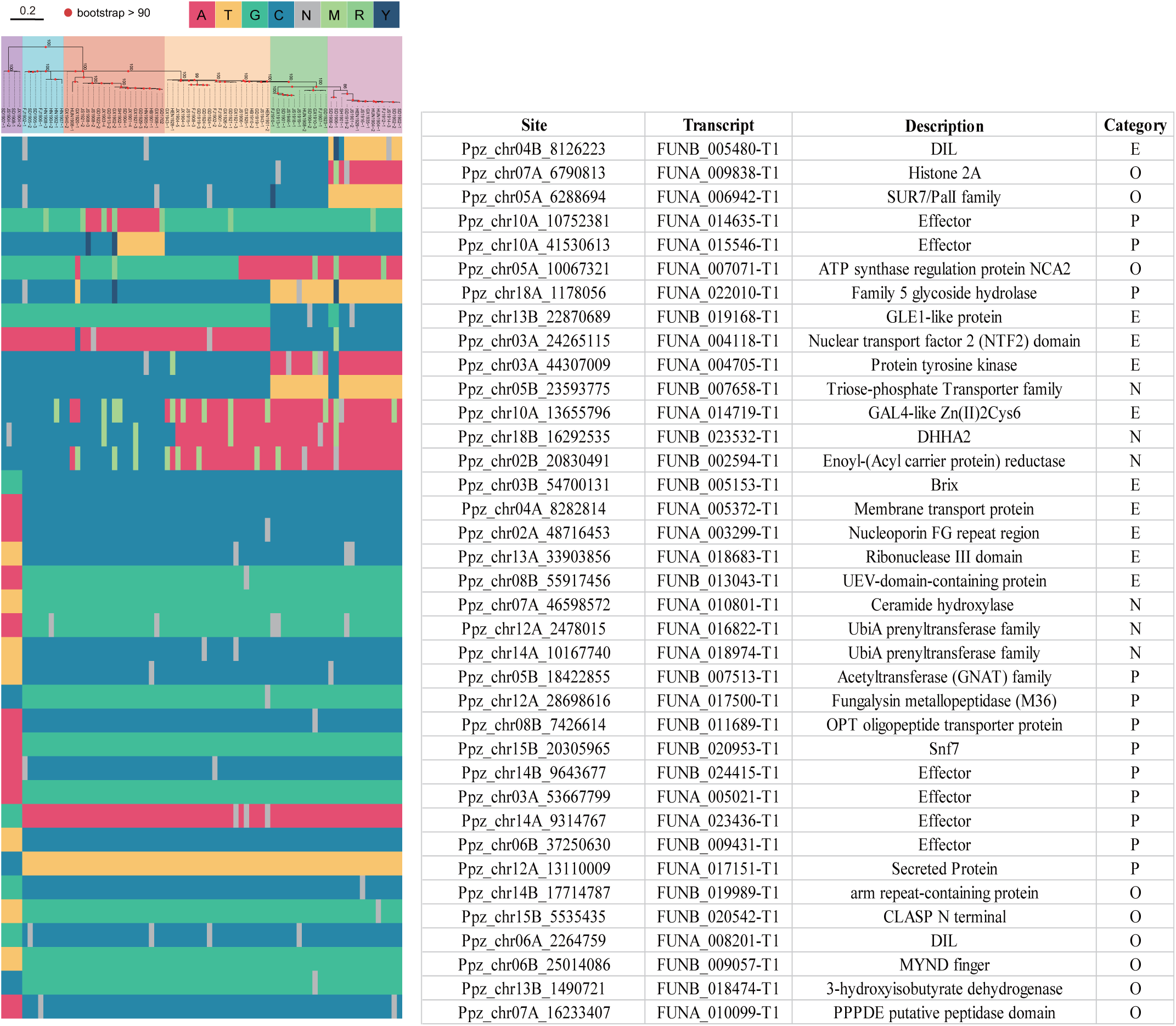
| Outlier SNPs (*Fst* > 0.9) and functional annotation of related genes in six groups. Schematic diagram showing outlier SNPs among six groups (left) and the corresponding gene function (right). Rows correspond to highly variable SNPs in each isolate (*Fst* > 0.9), and SNP allelic genotypes have been coloured according to the IUPAC Dictionary of ambiguities for nucleotides. Gene designations are based on the GD1913 *P. polysora* genome [27]. Category refers to COG category https://www.ncbi.nlm.nih.gov/research/cog).

### High variability of the effector repertoire in *P. polysora* populations

Given the inoculation phenotypic and genetic differentiation among groups that underscores the dynamic nature of *P. polysora* virulence, we assessed the effector repertoire on a population scale, aiming to provide comprehensive understanding of the virulence traits of *P. polysora.* By scanning complete open reading frame (ORF) of GD1913, 448 effectors were further captured (Fig. 5a) in addition to over thousand effectors based on gene structure in Liang et al (2023). Most effectors were shared between two haplotypes, with approximately 10.0% to 16.8% specific to hapA and hapB, respectively (Fig. 5b). Among all predicted effectors, 212 of them had low level of expression during the germination phase and early infection stages (1 dpi and 2 dpi), but showed a marked increase in expression by 4 dpi and 7 dpi (Fig. 5c, Table S4), suggesting that they are potential candidate avirulent effectors.

**Fig. 5.**
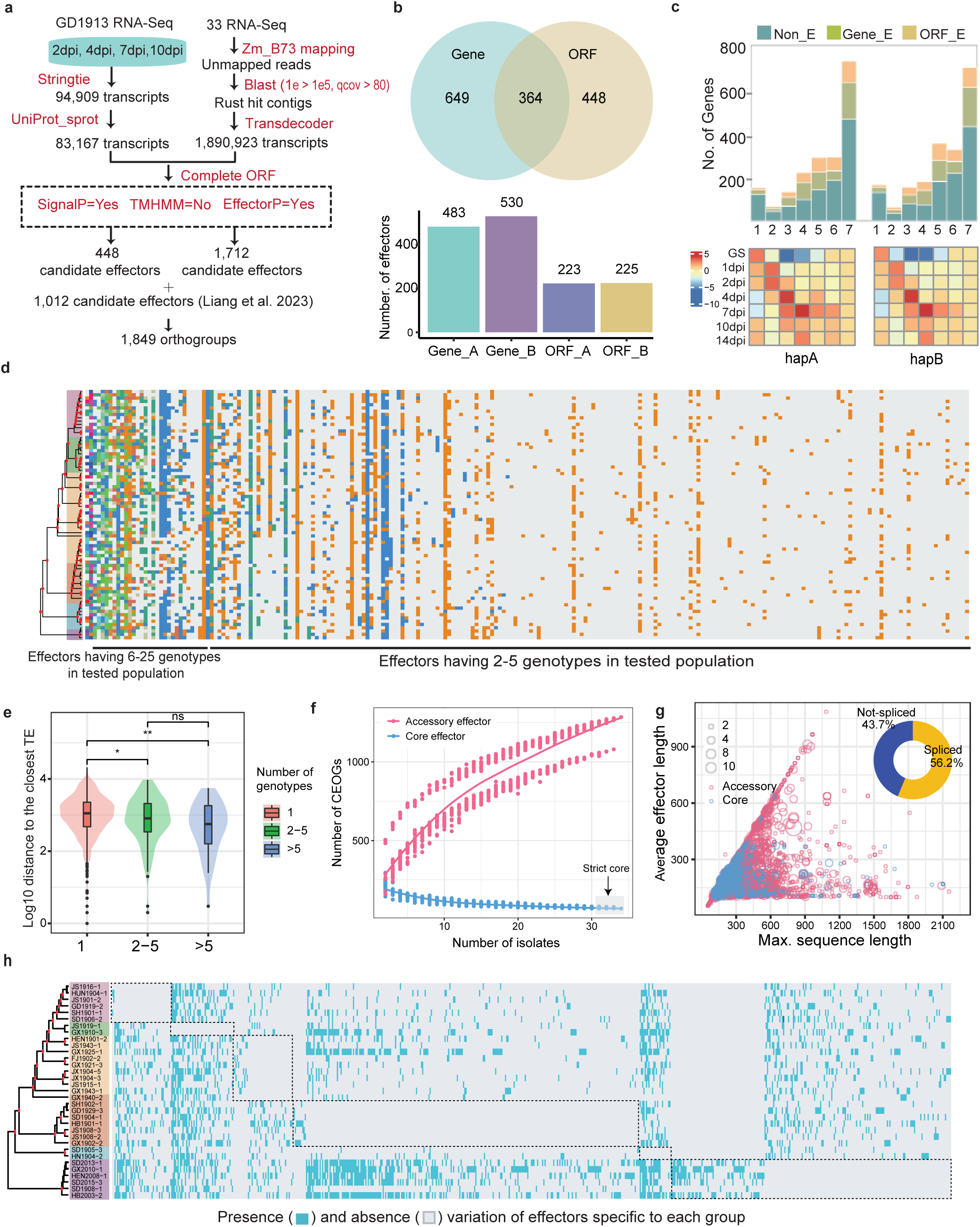
| Pan-effectorome landscape of *P. polysora* population. (a) Workflow to retrieve extra effectors ORF capture. (b) The Venn diagram of candidate effector orthogroups (CEOGs) predicted from two haplotypes of GD1913. (c) The top bar plot showing proportion of non-effector, effector in genome annotation and new effectors retrieved by ORF prediction from GD1913 in each expression cluster. The bottom heatmap showed different expression patterns of secreted protein at each time point of infection. (c) The top bar plot showing proportion of non-effector, effector in genome annotation and new effectors retrieved by ORF prediction in each expression cluster. The bottom heatmap showed different expression patterns of secreted protein at each time point of infection using rlog transformed expression values ranging from −10 to 5. The details of candidate effectors in each expression pattern were listed in Table S4. (d) Genotypic landscape of effectorome in *P. polysora* population. The ML tree shows the phylogenetic relationship of six groups. Different SNP types of all isolates (row) of each effector (column) are labelled in distinct colours. Gray colour means the same genotype compared to reference genome. For clarity, only effectors with more than two genotypes were visualized. (e) The log distances between effectors and their closest TE. The effectors were categorized as single genotype, 2-5 genotypes and > 5 genotypes across the population of 76 isolates. (f) Pan-effectorome accumulation curve illustrating core and accessory CEOGs with increasing isolate numbers. Core CEOGs were highlighted in grey box and listed in Table S7. (g) Protein sequence length of effectors and the corresponding longest transcript. Circle size indicates number of transcripts encoded by a effector. Pie chart shows the ratio of effectors via alternative splicing. (h) Presence/absence variation (PAV) of CEOGs in 33 *P. polysora* isolates with available RNA-seq data. The dashed boxes indicate CEOGs with PAV variation specific to each group.

We further analysed short variants of *P. polysora* population on all predicted effectors. A total of 367 variants (294 SNPs and 73 INDELs) were detected (Table S5). Approximately 40% of variants were rare, with the variants existed in a single isolate, resulting in dimorphic effectors in population. However, 25 effectors displayed high variability, presenting with 6-25 genotypes across populations (Fig. 5d, Table S5). Consistently, effectors with multiple genotypes exhibited significantly greater proximity to their neighbouring TEs than those with single genotype (Fig. 5e). The TE expression revealed that out of 25 highly variable effectors, 21 effectors have activated TEs (TPM > 0) around them, suggesting the potential impact of TEs on effector variants (Fig. S3).

We built the pan-effectorome to characterize effector repertoire of *P. polysora* population. The integration of transcriptomic data from 33 different isolates broadened the *P. polysora* effectorome and revealed 1,712 novel candidate effectors (Fig. 5a). On average, 388 effectors from 3.48% of assembled transcripts of each isolate were finally predicted (Table S6). Combined with the effectors from GD1913, a total of 3,173 non-redundant effectors were grouped into 1,849 candidate effector orthogroups (CEOGs) (Table S7).

To eliminate the effector diversity overestimation caused by insufficient expression, we further examined the effectors expression levels of each transcriptome data. Firstly, of the reference isolate GD1913, over 90% predicted effectors expressed at 14dpi, which was the same sampling timepoint as the 33 transcriptomic data (Table S8). Secondly, we examined representative effector CDS sequences of each CEOG within 33 transcriptomic data (Table S9). The strict and loose filter of CEOG expressed over 95% and 50% isolates, retained 1209 (65.39%) and 1466 (79.29%) CEOGs.

The core effectorome of *P. polysora* was significantly smaller, containing only 109 (5.9%) CEOGs (Fig. 5f, Table S9). The majority of CEOGs consisted of a single effector, highlighting the high effector diversity within this species. According to a protein sequences alignment test, the effectors that only one single of them grouped into one CEOG was 50% because genes with high similarity in other isolates were not predicted while other 50% contributed real effectorome diversity.

Among all effectors, alternatively spliced transcripts account for 56.2% (Fig. 5g). More importantly, 679 CEOGs are group-specific, suggesting that *P. polysora* employs a flexible accessory effector repertoire through differential gain/loss, thus rapidly generating a diverse array of genotypes with heterogeneous virulence profiles (Fig. 5h). Overall, *P. polysora* adopted a highly diversity effectorome to achieve adaptation.

### Mating-type loci and recombination

Successful mating in rust fungi typically requires the fusion of compatible cells, a process governed by genes that encode pheromone precursors (*mfa*) and their receptors (*STE3*), collectively referred to as the P/R locus, and heterodimeric transcription factors encoded by two homeodomain transcription factors, namely *bW-HD1* and *bE-HD2* [23]. To explore the genetic basis of sexual reproduction in *P. polysora*, we characterised the mating-type loci from the assembled genome. The *bW*/*bE* loci were anchored at ∼44.2 Mb and∼40.8 Mb in two haplotypes of chromosome 4, respectively (Fig. S4). For the STE loci, FUNA_013409 on chromosome 9, located near a *mfa*-like gene (FUNA_013420), showed 65%–75% identity to STE3.2-3 in other *Puccinia* species (Fig. 6a, Fig. S4). In hapB, a homologous gene, FUNB_023888, with 56%–67% identity to *STE3.2-2*, was found in an unplaced contig likely due to discontinuous assembly of GD1913 (Fig. 6a, Fig. S4). In addition, homologs of *STE3.2-1* were found as FUNA_001167 and FUNB_001172 on chromosome 1 with ∼64% identity (Fig. 6a, Fig. S4). The mating loci are generally conserved, except in G6, which has a SNP in the *bW* locus converting a tyrosine (Y) to histidine (H) (Fig. 6b). RNA-seq data confirmed the functional expression of these loci, revealing that *STE3.2-3* has a significantly higher expression level than *STE3.2-2* in most isolates, while *STE3.2-1* has the lowest expression level in hapA (Fig. S5).

**Fig. 6.**
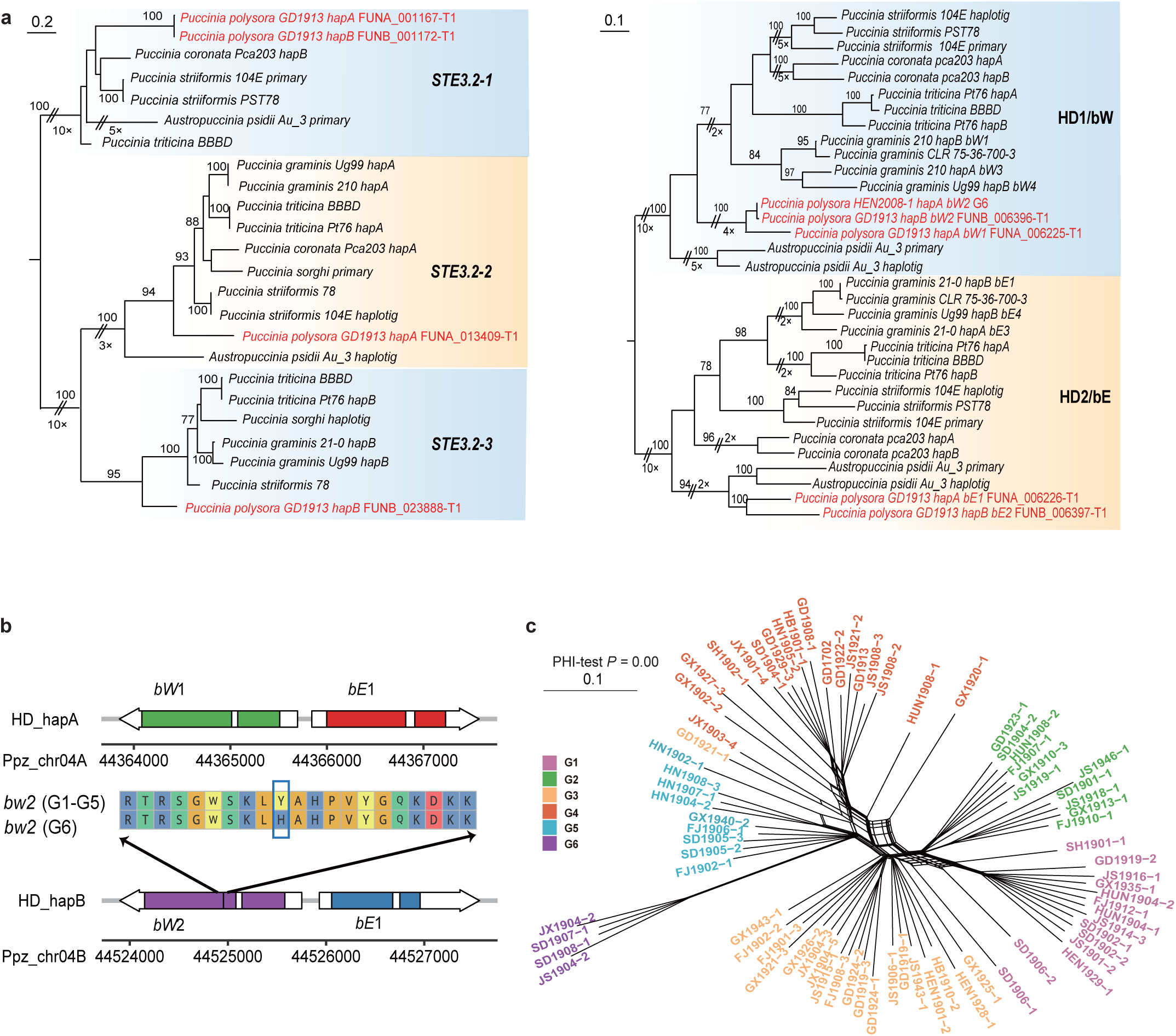
| Mating-type loci and recombination test of *Puccinia polysora*. (a) Phylogenetic trees constructed by mating-type loci of *P. polysora* and other rust fungi. (b) Gene structure of HD allele in tested population. The G6 specific non-synonymous variant on *bW*2 is marked by black bar. The region containing amino acid variation has been zoomed in. (c) Phylogenetic network analysis of *P. polysora* population. Network indicated recombination events and PHI-test *p-*value indicated recombination signal. Outlier isolates (e.g. HUN1908-1, GX1920-1, HN1902-1) contributed significant network.

The PHI-test revealed extensive recombination across the population, particularly in basal branches, with most isolates (especially those in G6) showing extended branches post network (Fig. 6c). The *K*-mer containment analysis revealed that two nuclei of all isolates have high similarity of two nuclei of GD1913 (Fig. S6b), excluding the possibility of somatic hybridization in contributing recombinational signal. Phylogenetic trees constructed using SNPs mapped to hapA and hapB showed different phylogenetic positions for SD1906-1 (G1), SD1906-2 (G1) and HN1902-1 (G5) (Fig. S6a). We further applied *K*-mer containment analysis to reveal nuclear differentiation of these three isolates in each chromosome. Larger internuclear variation were detected between two nuclei in these three isolates, for example, HN1902-1, SD1906-1 and SD1906-2 on chromosomes of Chr03, Chr07, and Chr110 (Fig. 7a). Phylogenetic trees using SNPs from each chromosome on both haplotypes were constructed and Robinson-Foulds distances of chromosome-pair trees suggested that Chr08, Chr11, Chr12, Chr13, Chr14, Chr 15, Chr 16 and Chr18 caused significant conflicts of two haplotype-based trees (Fig. 7b, Fig. S8). Especially in the tree of Chr13, all isolates in G5 have phylogenetic position shift (Fig. 7c). These potential internuclear exchanges may contribute to recombinational signal.

**Fig. 7.**
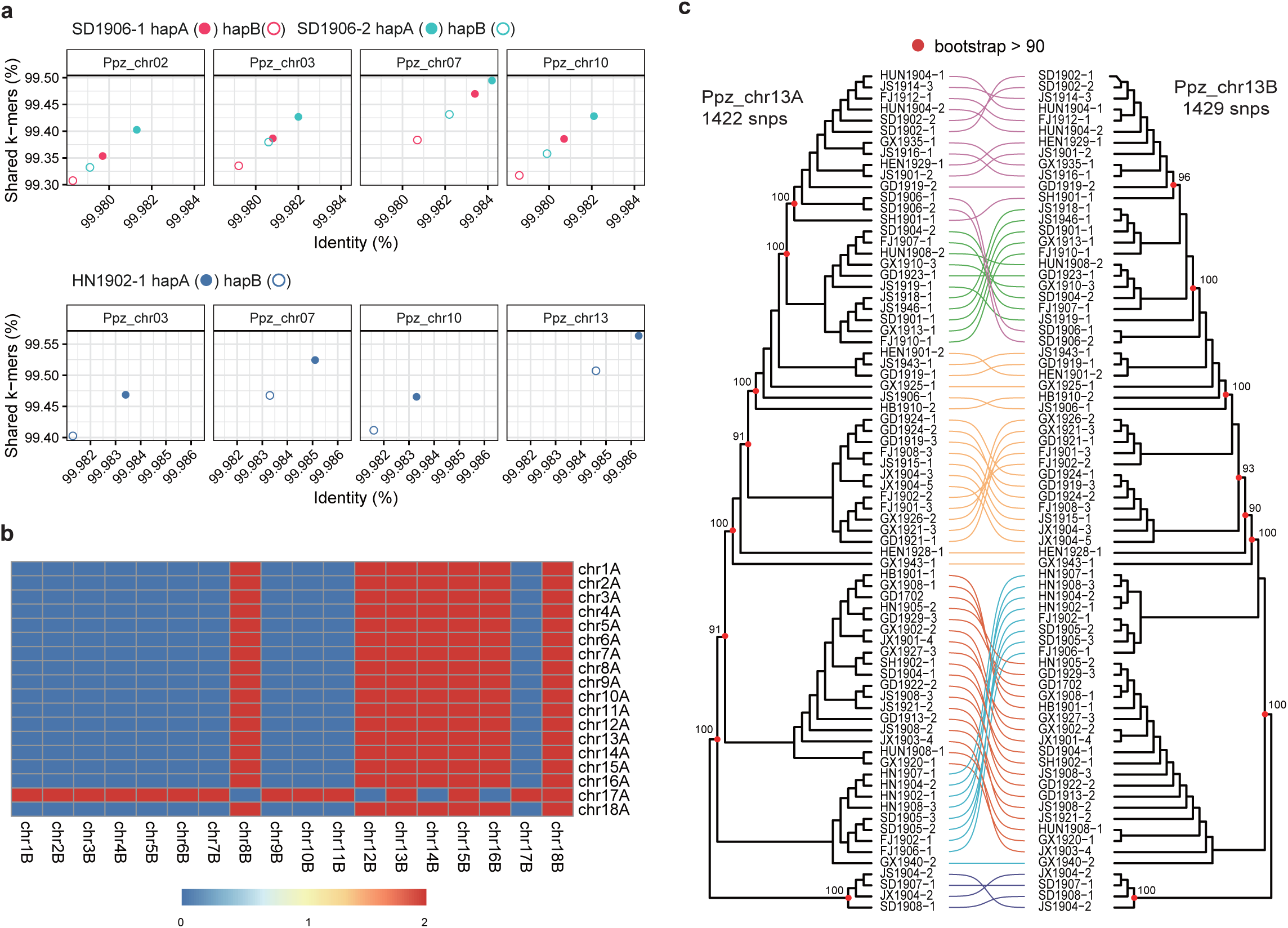
| (a) *K*-mer containment analyses of three outlier isolates against GD1913 chromosomes. X-axis stands for *K*-mer identity and Y-axis stands for percentage of shared *K*-mers (b) Robinson-Foulds distances between chromosomes on two haplotypes. Red colour stands for significant differences in group level tree topology between chromosomes. (c) The topology comparison of phylogenetic trees on chromosome Ppz_chr13.

## Discussion

*Puccinia polysora*, the causal agent of southern corn rust, is a fungal pathogen of global concern, due to its escalating threat to food security in the context of global warming. Our research came at the right time for a comprehensive study of this pathogen. We unravelled the unexpected genetic differentiation of the Chinese *P. polysora* population and highlighted the emergence of a highly virulent group. Through a comprehensive whole-genome scan and examination of the pan-effectorome, we pinpointed genes critical for driving group differentiation, shedding light on the genomic basis of virulence-driven population diversification. By analysing of mating gene diversity, phylogenetic network analysis and examining haplotype-specific SNP topologies, we elucidated the potential impact of internuclear exchange on the population differentiation. In addition, we demonstrated the importance of two haplotype sequences for accurate resequencing reads mapping and variant calling in dikaryotic fungi.

### The emergence of a highly virulent group

Our study revealed six distinct groups within the Chinese *P. polysora* population, in contrast to previously reported weak population differentiation [17]. These findings provide further evidence that genome-wide SNP analyses are more capable of uncovering previously unseen genetic groups [53,54]. In general, these groups do not correlate with their geographical origin, which may be due to the high capacity of rust spores to migrate and spread over long distances [55]. Wang et al. [56] had suggested that Chinese (mainland) populations of *P. polysora* has multiple origins and may be influenced by pathogens from the Philippines or Taiwan, China. In addition, China underwent five transitions in its maize varieties between 1982 and 2020 [31], with varieties of different genetic backgrounds sown in different areas. As an obligate biotroph, the urediniospores of *P. polysora* are dependent on the presence of viable maize plants for their survival [57]. Thus, the prolonged lack of hosts during the winter months (November to March) in the vast northern and central regions of China renders the *P. polysora* population extraneous in these regions. The different maize germplasm grown in China may be a driving force for pathogen differentiation [58], as the host plant has a significant influence on pathogen evolution. Therefore, the genetic differentiation observed within the Chinese population may be the result of an interplay between exotic inoculum and local adaptation.

The inoculation array indicated that we had discovered a highly virulent group, G6, which significantly diverged from other groups. The increased virulence can be attributed to the presence of specific allele types, *AvrRppC*^A^ and *AvrRppC*^J^, which have evaded the detection by their corresponding resistance (*R*) genes [27,59]. Inbred lines carrying the *RppC* resistance gene were particularly susceptible to G6 isolates but remained resistant to other groups (Fig. 3d, Fig. S2). We also detected group-specific variants in transporter (e.g., effectors) and protein modification genes in G6. Recent studies have highlighted the pivotal role of PAV in the evolution of virulence [50–52]. According to the pan-effectorome landscape of *P. polysora*, the differential gains/losses of effectors underscore their significant influence on the virulence spectrum. The similar virulence pattern among G1-G5 groups is likely due to the common recognition of avirulent genes by *RppQ*, *Rpp25*, *RppD* and*RppCML496*/*RppC*, which are clustered on the short arm of chromosome 10 of the maize genome [60,61]. Future research should focus on identifying additional resistance genes and establishing connections between *P. polysora* genetic groups and resistance mechanisms.

### The open pan-effectorome of *P. polysora*

Highly open pan-effectoromes are often found in species that possess multiple virulent “weapons” and diverse evasion strategies for evading host immune recognition. In our comprehensive analyses of 33 transcriptomic data from isolates across six groups, we observed a continuous increase in the number of effectors as effectoromes from new isolates were added. This suggests an open pan-effectorome of *P. polysora*.

Additionally, we highlighted the pivotal roles of activated TEs, PAVs and alternative splicing in the diversification of effectors. TEs are known to exert a substantial influence on gene regulation and sequence evolution of effectors [62], and they also contribute to structural variation that can modify the repertoire of effectors [26]. TEs (∼90%) are abundant in the genome of *P. polysora* [27]. In align with RIP test in other rust species, the RIP mechanism is also absent in *P. polysora*. Although some studies have shown that rust species can adopt RNA-directed DNA methylation (RdDM) to silence TEs [63–65], we detected a TE expression peak at 1 dpi in GD1913 (Fig. S3a). This temporally expression pattern of TEs was also detected in another giga-scale rust fungus with extreme repetitiveness (*Phakopsora pachyrhizi*) [66]. It is particularly noteworthy that effectors that are in proximity to TEs have more genotypes in tested population (> 5 genotypes in Fig. 5d). The effectors having 6-25 genotypes all have activated TEs (TPM > 0) nearby (Fig. S3b). This indicates a significant role for TEs in the evolution of virulence. In addition, PAV variants significantly expanded the effector diversity in *P. polysora*, as observed in *P. striiformis* [67], *Magnaporthe oryzae* [52], and *Zymoseptoria tritici* [51]. Copy number variants (CNVs), a proven key driver of genetic diversity in several fungal pathogens such as those causing powdery mildew [68], wheat stripe rust [69] and wheat leaf blotch [70], appeared to be absent in the *P. polysora* population, and no group-specific CNV patterns were detected (Fig. S9).

Alternative splicing is a process that enhances coding capacity in eukaryotes, but is less common in fungi than in animals and plants [71]. However, recent studies have highlighted its importance for fungal survival and virulence. For example, a non-effector gene in the cucurbit downy mildew pathogen was converted into a functional effector protein through alternative splicing [72], and a signal-specific isoforms determines the full virulence of *Magnaporthe oryzae* [73]. In our analysis, alternative splicing contributed more than half of the effectors in the *P. polysora* population (Fig. 5g). Despite excluding biases caused by different expression levels using transcriptomic data, the pan-effectorme of *P. polysora* remains open. This contrasts with our initial hypothesis that a clonal population would exhibit a homogeneous genetic structure, resulting in a closed pan-effectorome. We found that many ortholgous genes are expressed as different transcripts across strains. Some of them are no longer effectors due to factors such as the loss of signal peptides at the beginning or the addition of transmembrane structures. Therefore, the number of effectors ultimately obtained by each strain varied (Table S6), highlighting the significant impact of alternative splicing on the effectorome diversity. This may be an adaptive strategy for rust fungi to maintain rapid evolution during long-term asexual reproduction.

### Pitfalls and riches in genome analysis of dikaryotic fungi

Advances in sequencing technologies have improved our understanding of the genomes of rust fungi. Previous studies have primarily utilized collapsed genomes or monokaryotic genomes after nuclear phasing as references [74]. This approach artificially incorporates a large number of heterozygous sites or inter-nuclear differences into population polymorphisms [75], resulting in dilution of variation among isolates. Our approach maps short reads to both nuclei simultaneously, thereby accurately distinguishing between reads specific to either nucleus A or nucleus B. By eliminating the noise caused by SNPs between the two nuclei, we were able to uncover the true population structure of *P. polysora*. Phased genomes allow for in-depth analysis of variations within each nucleus, thereby addressing challenges associated with population-level nuclear exchange and structural variation, as discussed by Guo et al. [26] and Sperschneider et al. [28]. Our research serves as an illustrative example of strategy for mapping population genetics resequencing data of rust fungi, and provides a valuable case study for future investigations in this field.

### Recombination in *P. polysora*

The life cycle of *P. polysora* was an enigma, with uncertainties about its capacity for sexual reproduction on alternative hosts. In the case of *P. striiformis*, the causal agent of wheat stripe rust, comparative studies have indicated that the formation of telia is less frequent in clonal populations than in those that engage in sexual reproduction [76]. In contrast, in *P. polysora*, telia were rarely observed in previous studies [38,39], and were never found in current field sampling in China. Adhering to the null hypothesis of sexual reproduction of rust life cycles [77], the production of telia may have been an ancestral trait of rust fungi that may have been lost in *P. polysora*. However, this hypothesis awaits confirmation by conclusive evidence of the full life cycle in other populations.

Recombination suppression in sexual chromosomes/mating type loci is known in both Ascomycota and Basidiomycota [78,79]. This mechanism is instrumental in promoting compatibility under inbreeding conditions and assists in the integration of genes associated with sexual reproduction into mating type loci. Our results suggest that consistent with other rust species, the pheromone receptor (at the *STE* loci) reveals recombination suppression. *STE3.2-2* and *STE3.2-3*, two compatible alleles are highly conserved in *P. polysora*. However, the impact of recombination suppression in the *HD* locus (*bW*-*HD*1 and *bE*-*HD*2) is difficult to assess based on current available data. Usually, the HD loci include three structures: N-terminal Variable domain at the N-terminal, a Homeodomain and a Constant domain at C-terminal. In *P. striiformis*, at least nine *bW* alleles showing variation at N-terminal have been detected in the global population, particularly with high diversity in the populations that sexual reproduction occurs [42]. In our study, *bE* is conserved but a single SNP in the Homeodomain on *bW* detected in G6. Future detection on HD loci in other regional populations will be helpful to answer this question thoroughly. Nevertheless, the low diversity of *bW*-*HD*1 and *bE*-*HD*2 suggested a predominantly clonal nature of the Chinese *P. polysora*.

In the absence of sexual reproduction, mutation, internuclear exchange and somatic hybridisations are three hypothesized modes that drive virulence evolution in rust species during asexual reproduction [74]. Somatic hybridisation has recently been validated in *P. graminis*. Its highly virulent race, Ug99, was identified to have a completely different nucleus compared to a dominant race [24,80]. Additionally, extensive nuclear exchange events have been detected in the global populations of *P. triticina* [28] and *P. coronata* [81]. However, the *K*-mer analysis suggested the high similarity of the nuclear of all isolates to two phased isolates of GD1913, excluding the possibility of somatic hybridisation in *P. polysora* population. However, we found some outliers, such as SD1906-1, SD1906-2 and HN1902-1, their phylogenetic positions varied between two haplotypes on chromosomes 13, 16 and 18. The SNPs on chromosome 13 even impacted the phylogenetic positions of all isolates in G5. These conflicts in the topology of phylogenetic trees based on two nuclei imply internuclear exchange existed in *P. polysora* population. Our study provides a field case to explain how internuclear exchange plays roles in admixing events in the clonal population.

## Materials and Methods

### Isolate collection, culturing and inoculation assay

In this work, we sequenced an additional new isolate (GD1702) and used 75 genome resequencing data (Table S10) from our previous work [27] to reassess the population genetic structure of *P. polysora*. We also constructed RNA-seq from the 28 isolates above and an additional five isolates from a highly diverged group to look for signatures of adaptation. These isolates were collected from epidemic regions in China (Fig. 1b) and propagated from a single pustule according to the protocol described by Liang et al (2023) in green house. The purified isolates were inoculated onto the second leaf of the susceptible cultivar, Zhengdan 958 and underwent 2-3 rounds of amplification. At 14 days post inoculation (dpi), intensively infected leaves (∼5 cm in length) were cut down and frozen in liquid nitrogen for RNA extraction. For DNA extraction, the single pustule was amplified by 3-4 times to obtain sufficient spores (∼10 mg).

To demonstrate the differentiation of virulence among different groups, 12 representative isolates were inoculated onto five inbred lines carrying known resistance genes, namely CML470 (*RppC*) [82], CML496 (*RppCML496, RppC*) [59], F939 (*RppP25*) [83], W2D (*RppD*) [84] and Qi319 (*RppQ*) [85]. The inbred lines were provided by maize breeding teams from the State Key Laboratory of Plant Environmental Resilience, China Agricultural University and Tianjin Academy of Agricultural Sciences. The seedlings of the above inbred lines were sown 7-10 days before inoculation, and inoculated together with 10 mg urediniospores suspended in 15 mL of Tween suspension (0.05%). Incubation procedures were described by Liang et al. [27]. The susceptible variety Zhengdan 958 was inoculated as a positive control for three biological replicates.

### RNA/DNA extraction and sequencing

Total RNA was extracted using the TRIzol reagent (Invitrogen, Carlsbad, CA, USA), and DNA was extracted using QIAGEN kits (DNeasy Plant Mini Kit, QIAcube HT). Nucleotide quality was assessed using an Agilent 4200 Bioanalyzer (Agilent Technologies, CA, USA). Sequencing libraries with a size of approximately 350 bp were prepared from sheared RNA/DNA. Libraries were sequenced on the Illumina NovaSeq 6000 platform, generating 150 bp pair-end reads at Annoroad Gene Technology Co., Ltd. (Beijing, China).

### Read mapping and variant calling

For population genomic analyses, the short reads were trimmed using the settings of “*ILLUMINICLIP 2:30:10 LEADING 3, TRAILING 3, SLIDINGWINDOW 4:10, MINLEN 50*” in Trimmomatic v 0.36 [86] and then aligned to hapA+hapB (tandem mapping mode) or hapA/hapB (single haplotype) using HISAT2 v 2.2.1 [87] with --no-spliced-aglignment for genomic data mapping and others as default settings. The resulting SAM files were sorted and converted to BAM format via SAMtools v1.9 [88]. Finally, variants were retrieved using Freebayes v 1.3.2 [89] and filtered using vcflib v 1.0.0 [90] with the following parameters: QUAL > 20 & QUAL/AO > 10 & SAF > 0 & SAR > 0 & RPR > 1 & RPL > 1 & AC > 0 & FS < 60. Only the biallelic SNPs with depth > 5, minor allele frequency (MAF) > 0.05 and missing data < 90% were retained for phylogenetic analysis. The effect (e.g. non-synonymous or synonymous) and related genes of the variants were annotated using SnpEff v 4.3 [91]. The maximum likelihood (ML) tree was constructed by loading the full set of quality-filtered SNPs into IQ-Tree [92] and visualized in the R package ggtree v.3.2.1 [93]. The best-fit evolutionary model was determined according to BIC using -m MFP+ASC. The phylogenetic relationships of the *P. polysora* population were compared between the two mapping strategies. The heterozygosity of each isolate was determined by using vcftools -het [88].

### Population genomic analyses

We used the filtered biallelic SNPs to estimate the population structure of *P. polysora*. The discriminant analysis of principal components (DAPC), a model-free K-means clustering methodwas performed using R package adegenet v 2.0.1 [94] with genetic clusters (*K*) ranging from 2 to 7. The optimal *K* value was determined by the lowest Bayesian Information Criterion (BIC) value generated in find.cluster() function. Nucleotide diversity (*Pi*) and pair-wise fixation index (*F*st) were calculated using vcftools within a non-overlapping 100 Kb window [95]. We concentrated the outlier variants with high *F*st (> 0.9) between groups. Principal component analysis (PCA) was performed using PLINK v1.9 [96] and visualized using ggplot [97]. In addition, to test for the presence of repeat-induced point mutation (RIP), two dinucleotide ratios TpA/ApT and (CpA + TpG)/(ApC+GpT) were calculated for each type of TEs in RIPCAL v2.0.0 [98]. Repeat positions followed the prediction in the *P. polysora* reference genome, GD1913 [27].

### Pan-effectorome construction

In order to comprehensively understand the effector diversity of *P. polysora* at the population level, we assessed SNP variation based on effectors predicted from the reference genome, GD1913, as well as presence/absence variation based on RNA-seq data from 33 representatives of each group. Our previous work [27] has predicted over a thousand candidate effectors in GD1913. However, a recently identified avirulent effector, *AvrRppC*, was not included due to its incomplete gene structure and was not annotated as a gene (Deng et al., 2022; Liang et al., 2023). Therefore, we expanded our approach by scanning the complete open reading frame (ORF) to capture those unannotated effectors in GD1913.We used Stringtie v 2.2.1 [99] to merge all transcripts derived from the infection transcriptome data of 2 dpi, 4 dpi, 7 dpi and 10 dpi from Liang et al. [27]. And the ORFs were then identified using Transdecoder v 5.5.0 [100]. Only the longest and labeled as complete (containing both start and stop codons) were filtered by SignalP v 4.1 [101], TMHMM v 2.0 [102] and EffectorP v 3.0 [103]. The expression patterns of all effectors predicted from GD1913 were analysed according to the protocol described by Liang et al. [27]. To uncover presence/absence or pan-effectorome of *P. polysora* population, we scanned the ORF of 33 randomly selected isolates from six groups (Table S10). For a given group (G6), five additional isolates sampled at 14 dpi were supplemented due to only one isolate with available RNA-seq. The phylogenetic tree based on RNA-seq suggested that these five isolates clustered together with the one in G6 (Fig. S11). RNA-seq reads were first mapped to the maize genome B73 v5[104] to filter host sequence. The unmapped ORFs were extracted by Transdecoder v 5.5.0 [100] following the steps performed in GD1913. Then the transcripts were remained by following three criteria: 1) length longer than 200 bp, 2) with ‘complete’ flag in the transcript title, 3) hit to any rust genera of Pucciniales in nr database (e > 1e-5, qcovs > 80). The detail codes are available at https://github.com/cpwater/PopGenomePPolysora. To show the presence/absence variation of effectors among different groups, all predicted effectors were grouped to Candidate Effector Orthogroups (CEOG) using Orthofinder v 2.5.4 with parameters ‘-M msa -S diamond’ and diamond parameters were set to ‘-e 1e-5 - i 80’ [105]. To reduce the bias due to expression levels, only the CEOG expressed in over 95% of isolates were used for statistics.

TE expression was calculated using TEtranscript [106]. The timecourse RNA-seq data sets were mapped to GD1913 reference genome using Hisat v 2.2.1 [87] as mentioned above and duplicated reads were removed using Picard v 2.23 (https://broadinstitute.github.io/picard/). The resulting data was used to generate counts in TE transcripts along with TE gtf file from GD1913 genome annotation files. TE reads counts were normalised to generate a CPM matrix in edgeR v 3.1 [107]. Expressed TEs were defined as with more than one read in at least 2 replicates of each time-point.

### Recombination test

We searched the mating-type loci of other *Puccinia* species (Table S11) against two haplotypes of *P. polysora* GD1913, respectively. The unrooted trees based on the homologous protein sequences of *STE3* and *bE*/*bW* were constructed using IQ-Tree [92]. The expression levels in transcripts per million (TPM) of each mating gene were analysed by mapping RNA-seq data to the phased genome in Kallisto v 0.48 [108].

To infer the recombination potential of *P. polysora* at the population level, a split network was constructed using the PHILIP sequence in the phylogenetic tree in SplitsTree v 4.0 [109] The pairwise homoplasy index (PHI) test was also performed to detect recombination using PhiPack [110]. To further access the linkage disequilibrium level of each group, we calculated the standardized association index (rd). The 100 sets of 10,000 random SNPs were constructed using the samp.ia function from the R package *poppr* to generate a distribution of rd values [111]. The observed rd distribution of each group was compared to the distribution of 10,000 rd values constructed using fully randomly simulated datasets with 0% (fully sexual), 25%, 50%, and 100% linkage levels. To detect the effects of somatic hybridization or internuclear exchange in *P. polysora*, we extracted the variants belonging to either hapA or hapB from the filtered vcf file mapping in tandem mode. These two subdatasets were used to construct two phylogenetic trees, which were visualized in ggtree v 3.2.1 [93]. K-mer containment analysis was further performed using Mash v2.3 [112]. After sketching sequences of each chromosomes using parameter -s 500000 -k 32, mash screen was run to get identity and shared *K*-mer for each isolates sequencing data. Phylogenetic trees on each chromosome were constructed. The topology differences between on two haplotypes were visualised using ggtree v 3.2.1 [93]. Robinson-Foulds (RF) distance was calculated using the R package ape v 5.8 [113] on subtrees consisting of representative isolates within each genetic group, with higher RF distance values indicating greater topological differences and thus more dissimilar evolutionary relationships.

## Data availability

Illumina sequence files for the *P. polysora* population are available from National Microbiology Data Centre (NMDC, https://nmdc.cn/) under the BioProject NMDC10018113.

## Acknowledgments

We would like to thank following persons for their assistance in collecting samples: people from Universities or research institutes, Keyu Zhang, Leifu Li, Shiling Han, Jun-en Huang, Qing Zuo, Rijian Wei, Zhaohong Liu, Yuying Lao, Yunhe Xu, Yanqi Zheng, Wei Wang, Hailong Er, Liangliang Wang and Yu Shi; persons from regional plant protection stations, Peng Lu, Gongxian Liao, Shikun Pan, Xingshang Lu, Baoqin Tan, Yanqi Zheng, and Xingren He. It’s also grateful to Prof. Zhanhong Ma and Prof. Guozhi Bi from China Agricultural University, for supplying inbred lines for inoculation assay. The work was financially supported by the Strategic Priority Research Program of Chinese Academy of Sciences (XDB0830000), National Sciences Foundation of China (NSFC 32330002, 32472506, 31972210) and the Key Collaborative Research Program of the Alliance of International Science Organizations (ANSO-CR-KP-2022-07).

## Author contributions

J.M.L and L.C conceived the study; Y.J.L. and J.M.L carried out the experiment and performed analyses; X.F.L contributed samples and inbred lines. The manuscript was primarily written by J.M.L and Y.J.L, with critical input and revisions from L.C., C.K.M.T., and D.C.

